# Notch and TLR signaling coordinate monocyte cell fate and inflammation

**DOI:** 10.1101/2020.04.28.066217

**Authors:** Jaba Gamrekelashvili, Tamar Kapanadze, Stefan Sablotny, Corina Ratiu, Khaled Dastagir, Matthias Lochner, Susanne Karbach, Philip Wenzel, Andre Sitnow, Susanne Fleig, Tim Sparwasser, Ulrich Kalinke, Bernhard Holzmann, Hermann Haller, Florian P. Limbourg

## Abstract

Conventional Ly6C^hi^ monocytes have developmental plasticity for a spectrum of differentiated phagocytes. Here we show, using conditional deletion strategies in a mouse model of Toll-like receptor (TLR) 7-induced inflammation, that the spectrum of developmental cell fates of Ly6C^hi^ monocytes, and the resultant inflammation, is coordinately regulated by TLR and Notch signaling. Cell-intrinsic Notch2 and TLR7-Myd88 pathways independently and synergistically promote Ly6C^lo^ patrolling monocyte development from Ly6C^hi^ monocytes under inflammatory conditions, while impairment in either signaling axis impairs Ly6C^lo^ monocyte development. At the same time, TLR7 stimulation in the absence of functional Notch2 signaling promotes resident tissue macrophage gene expression signatures in monocytes and ectopic differentiation of Ly6C^hi^ monocytes into macrophages and dendritic cells, which appear in the blood stream and infiltrate the spleen and major blood vessels, resulting in unrestrained systemic inflammation. Thus, Notch2 is a master regulator of Ly6C^hi^ monocyte cell fate and inflammation in response to TLR signaling.

## Introduction

Infectious agents or tissue injury trigger an inflammatory response that aims to eliminate the inciting stressor and restore internal homeostasis (J. Bonnardel & Guilliams, 2018). The mononuclear phagocyte system (MPS) is an integral part of the inflammatory response and consists of the lineage of monocytes and macrophages (MF) and related tissue-resident cells. A key constituent of this system are monocytes of the major (classic) monocyte subtype, in mice called Ly6C^hi^ monocytes. They originate from progenitor cells in the bone marrow (BM), circulate in peripheral blood (PB) and respond dynamically to changing conditions by differentiation into a spectrum of at least three distinct MPS effector phagocytes: MF, dendritic cells (DC), and monocytes with patrolling behavior (Arazi et al., 2019; J. Bonnardel & Guilliams, 2018; Chakarov et al., 2019; Gamrekelashvili et al., 2016; Hettinger et al., 2013). The diversity of monocyte differentiation responses is thought to be influenced by environmental signals as well as tissue-derived and cell-autonomous signaling mechanisms to ensure context-specific response patterns of the MPS (Okabe & Medzhitov, 2016). However, the precise mechanisms underlying monocyte cell fate decisions under inflammatory conditions are still not fully understood.

When recruited to inflamed or injured tissues, Ly6C^hi^ monocytes differentiate into MF or DC with a variety of phenotypes and function in a context-dependent-manner and regulate the inflammatory response (Krishnasamy et al., 2017; Xue et al., 2014). However, Ly6C^hi^ monocytes can also convert to a second, minor subpopulation of monocytes with blood vessel patrolling behavior, in mice called Ly6C^lo^ monocytes (Gamrekelashvili et al., 2016; Patel et al., 2017; Varol et al., 2007; Yona et al., 2013). These monocytes have a long lifespan and remain mostly within blood vessels, where they crawl along the luminal side of blood vessels to monitor endothelial integrity and to orchestrate endothelial repair (Auffray et al., 2007; Carlin et al., 2013; Getzin et al., 2018). Steady-state monocyte conversion occurs in specialized endothelial niches and is regulated by monocyte Notch2 signaling activated by endothelial Notch ligands (Avraham-Davidi et al., 2013; Bianchini et al., 2019; Gamrekelashvili et al., 2016; Varol et al., 2007). Notch signaling is a cell contact-dependent signaling pathway regulating cell fate decisions in the innate immune system (Radtke, Macdonald, & Tacchini-Cottier, 2013). Notch signaling regulates formation of intestinal CD11c^+^CX3CR1^+^ immune cells (Ishifune et al., 2014), Kupffer cells (Johnny Bonnardel et al., 2019; Sakai et al., 2019) and macrophage differentiation from Ly6C^hi^ monocytes in ischemia (Krishnasamy et al., 2017), but also development of conventional DCs (Caton, Smith-Raska, & Reizis, 2007; Epelman et al., 2014; Lewis et al., 2011), which is mediated by Notch2.

Toll-like receptor 7 (TLR7) is a member of the family of pathogen sensors expressed on myeloid cells. Originally identified as recognizing imidazoquinoline derivatives such as Imiquimod (R837) and Resiquimod (R848), TLR7 senses ssRNA, and immune-complexes containing nucleic acids, in a Myd88-dependent manner during virus defense, but is also implicated in tissue-damage recognition and autoimmune disorders (Kawai & Akira, 2010). TLR7-stimulation induces cytokine-production in both mouse and human patrolling monocytes and mediates sensing and disposal of damaged endothelial cells by Ly6C^lo^ monocytes (Carlin et al., 2013; Cros et al., 2010), while chronic TLR7-stimulation drives differentiation of Ly6C^hi^ monocytes into specialized macrophages and anemia development (Akilesh et al., 2019). Furthermore, systemic stimulation with TLR7 agonists induces progressive phenotypic changes in Ly6C^hi^ monocytes consistent with conversion to Ly6C^lo^ monocytes, suggesting involvement of TLR7 in monocyte conversion (M.-L. Santiago-Raber et al., 2011). Here, we show that Notch signaling restrains TLR-driven inflammation by modulating Ly6C^lo^ monocyte vs. macrophage cell fate decisions in inflammation.

## Results

### TLR and Notch signaling promote monocyte conversion

We first studied the effects of TLR and/or Notch stimulation on monocyte conversion in a defined *in vitro* system (Gamrekelashvili et al., 2016). Ly6C^hi^ monocytes isolated from the bone marrow of *Cx3cr1^gfp/+^* reporter mice (GFP^+^) were cultured with recombinant Notch ligand DLL1 in the presence or absence of the TLR7/8 agonist R848 and analyzed after 24 hours for the acquisition of key features of Ly6C^lo^ monocytes (Gamrekelashvili et al., 2016; Hettinger et al., 2013). In contrast to control conditions, cells cultured with DLL1 showed an upregulation of CD11c and CD43, remained mostly MHC-II negative, and expressed transcription factors *Nr4a1* and *Pou2f2,* leading to a significant, five-fold increase of Ly6C^lo^ cells, consistent with enhanced monocyte conversion. Cells cultured with R848 alone showed a comparable phenotype response, both qualitatively and quantitatively (Figure 1A-C). Interestingly, on a molecular level, R848 stimulation primarily acted on *Pou2f2* induction, while Notch stimulation primarily induced *Nr4a1*. Furthermore, the combination of DLL1 and R848 strongly and significantly increased the number of CD11c^+^ CD43^+^ Ly6C^lo^ cells above the level of individual stimulation and significantly enhanced expression levels of both transcriptional regulators *Nr4a1* and *Pou2f2* (Figure 1A-C), suggesting synergistic regulation of monocyte conversion by TLR7/8 and Notch signaling. By comparison, the TLR4 ligand LPS also increased Ly6C^lo^ cell numbers and expression levels of *Nr4a1* and *Pou2f2*. However, the absolute conversion rate was lower under LPS and there was no synergy with DLL1 (Figure 1D and E).

**Figure 1.**
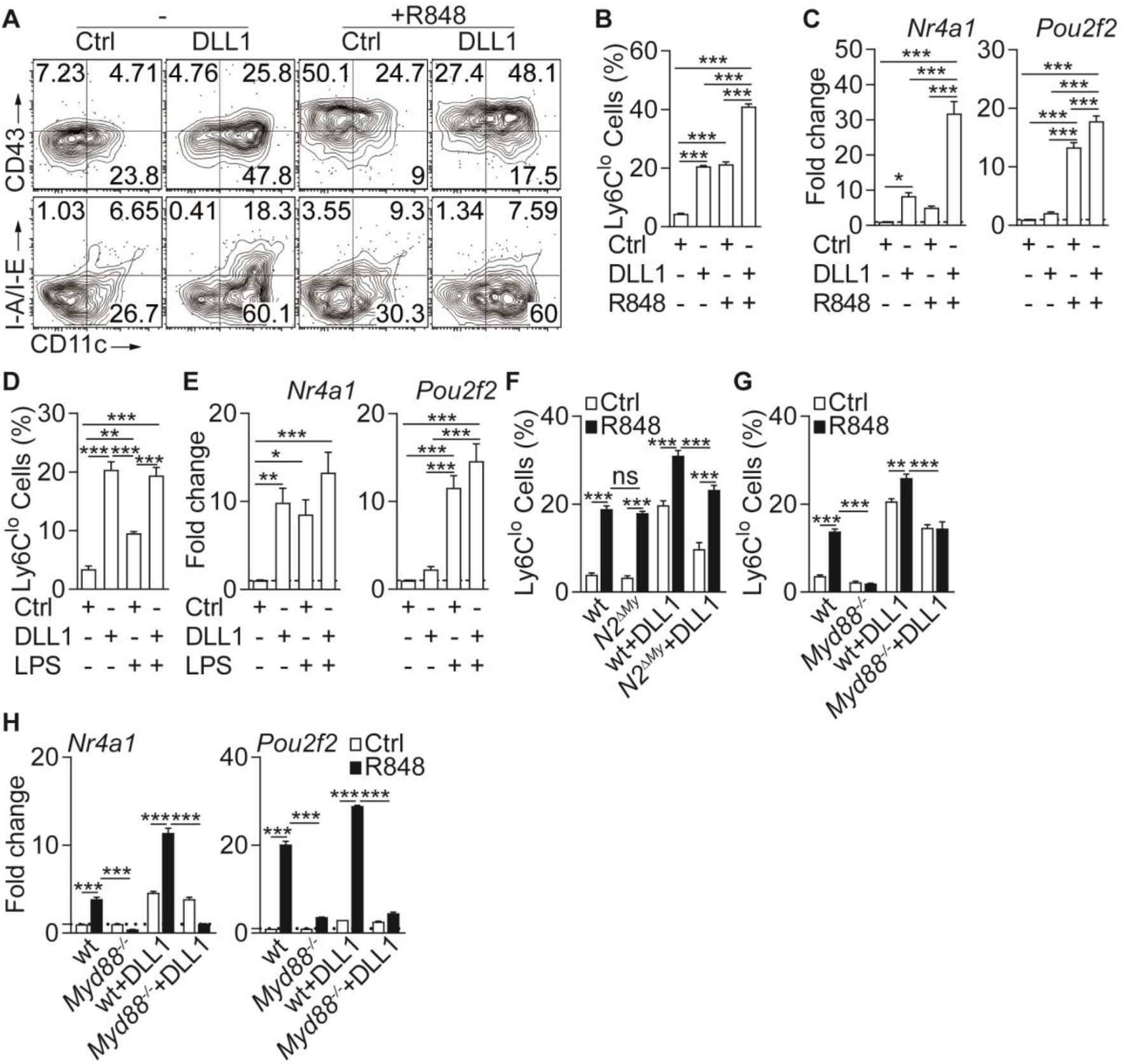
Inflammatory conditions enhance monocyte conversion *in vitro.* (**A-F**) Monocyte conversion in the presence of DLL1 and TLR agonists *in vitro:* (**A**) Representative flow cytometry plot, (**B**) relative frequency of Ly6C^lo^ monocyte-like cells (representative of 3 experiments, n=3) and (**C**) gene expression (pooled from 4 experiments, n=8-12) in the presence of R848 are shown. (**D**) Relative frequency of Ly6C^lo^ monocyte-like cells (from 3 experiments, n=3) and (**E**) expression of *Nr4a1* and *Pou2f2* (from 4 experiments, n=4-6) in the presence of LPS *in vitro* are shown. (**F**) wt or *N2^ΔMy^* Ly6C^hi^ monocyte conversion in the presence of DLL1 and R848 *in vitro:* relative frequency of Ly6C^lo^ monocytelike cells (from 3 experiments, n=4) is shown. (**G, H**) R848-enhanced conversion is *Myd88* dependent *in vitro*. Relative frequency of Ly6C^lo^ monocyte-like cells (**G**) and gene expression analysis *in vitro* (**H**) are shown (data are from two independent experiments, n=3). (**B-H**) * *P*<0.05, ** *P*<0.01, *** *P*<0.001; 2way ANOVA with Bonferroni’s multiple comparison test.

Since monocyte conversion is regulated by Notch2 *in vitro* and *in vivo* (Gamrekelashvili et al., 2016), we next tested TLR-induced conversion in Ly6C^hi^ monocytes with conditional deletion of myeloid Notch2 (*N2^ΔMy^*). Both, wt and *N2^ΔMy^* monocytes showed comparable response to R848, but conversion in the presence of DLL1, and importantly, also DLL1-R848 co-stimulation was significantly impaired in mutant cells (Figure 1F). This suggests independent contributions of TLR and Notch signaling to monocyte conversion.

To study whether the TLR stimulation requires Myd88 we next tested Ly6C^hi^ monocytes with Myd88 loss-of-function (*Myd88^−/−^*). Compared to wt cells, *Myd88^−/−^* monocytes showed strongly impaired conversion in response to R848 but a conserved response to DLL1. The response to DLL1-R848 co-stimulation, however, was significantly impaired (Figure 1G). Furthermore, expression of *Nr4a1* and *Pou2f2* by R848 was strongly reduced in *Myd88^−/−^* monocytes with or without DLL1 co-stimulation, while DLL1-dependent induction was preserved (Figure 1H). Thus, Notch and TLR signaling act independently and synergistically to promote monocyte conversion.

To address the role of TLR stimulation for monocyte conversion *in vivo* we adoptively transferred sorted Ly6C^hi^ monocytes from CD45.2^+^GFP^+^ mice into CD45.1^+^ congenic recipients, injected a single dose of R848 and analyzed transferred CD45.2^+^GFP^+^ cells in BM and Spl after two days (Figure 2A). Stimulation with R848 significantly promoted conversion into Ly6C^lo^ monocytes displaying the proto-typical Ly6C^lo^CD43^+^CD11c^+^MHC-II^lo/-^ phenotype (Figure 2B and C). In contrast, transfer of *Myd88^−/−^* Ly6C^hi^ monocytes resulted in impaired conversion in response to R848 challenge (Figure 2D and E). Together, these data indicate that TLR and Notch cooperate in the regulation of monocyte conversion.

**Figure 2.**
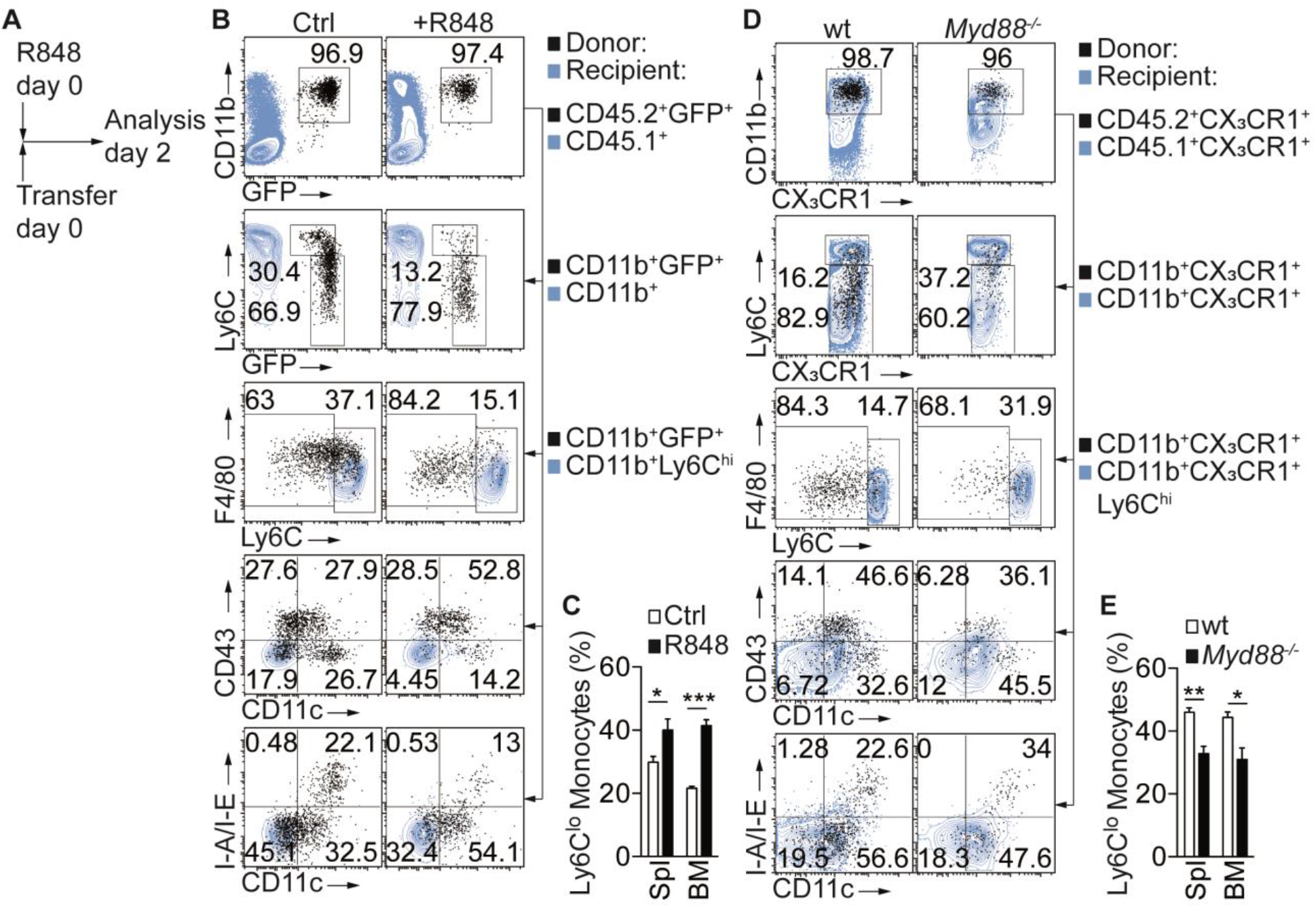
Inflammatory conditions enhance monocyte conversion *in vivo.* (**A-E**) Adoptive transfer and flow cytometry analysis of BM CD45.2^+^ Ly6C^hi^ monocytes in control or R848 injected CD45.1 ^+^ congenic recipients: (**A**) Experimental setup is depicted; (**B**) Flow cytometry plots showing transferred cells in black and recipient CD45.1^+^ (1^st^ row), CD45.1^+^CD11b^+^ (2^nd^ row) or CD45.1^+^CD11b^+^Ly6C^hi^ cells (3^rd^ −5^th^ rows) in blue; (**C**) Frequency of donor-derived LyGC^lo^ monocytes pooled from two independent experiments (n=5). (**D, E**) R848-enhanced conversion is *Myd88* dependent *in vivo.* (**D**) Flow cytometry plots showing transferred CD45.2^+^CX_3_CR1^+^ wt or *Myd88^−^’* cells in black and recipient CD45.1^+^CX_3_CR1 ^+^ (1^st^ row), CD45.1^+^CX_3_CR1^+^CD11b^+^ (2^nd^ row) or CD45.1^+^CX_3_CR1^+^CD11b^+^Ly6C^hi^ cells (3^rd^ −5^th^ rows) in blue; (**E**) Frequency of donor-derived Ly6C^lo^ monocytes pooled from two independent experiments are shown (n=4/5). (**C, E**) * *P*<0.05’ ** *P*<0.01, *** *P*<0.001; Student’s *t*-test.

### Notch2-deficient mice show altered myeloid inflammatory response

To characterize the response to TLR stimulation *in vivo* we applied the synthetic TLR7 agonist Imiquimod (IMQ, R837) in a commercially available crème formulation (Aldara) daily to the skin of mice (El Malki et al., 2013; van der Fits et al., 2009) and analyzed the systemic inflammatory response in control or *N2^ΔMy^* mice (Gamrekelashvili et al., 2016) (Figure 3A). While treatment with IMQ induced comparable transient weight loss and ear swelling in both genotypes (Figure S1A), splenomegaly in response to treatment was significantly more pronounced in *N2^ΔMy^* mice (Figure S1B).

To characterize the spectrum of myeloid cells in more detail we next performed flow cytometry of PB cells with a dedicated myeloid panel (Gamrekelashvili et al., 2016) and subjected live Lin^−^CD11b^+^GFP^+^ subsets to unsupervised t-SNE analysis (Figure 3B). This analysis strategy defined 5 different populations, based on single surface markers: Ly6C^+^, CD43^+^, MHC-II^+^, F4/80^hi^ and CD11c^hi^ (Figure 3C). Applying these 5 gates to samples from separate experimental conditions identified dynamic alterations in blood myeloid subsets in response to IMQ, but also alterations in *N2^ΔMy^* mice (Figure 3D). Specifically, abundance and distribution of Ly6C^+^ cells, containing classical monocytes, in response to IMQ were changed to the same extend in both genotypes. In contrast, the MHC-II^+^ and F4/80^hi^ subsets were more abundant in *N2^ΔMy^* mice, but also showed more robust changes in response to IMQ. On the other hand, the CD43^+^ subset, containing the patrolling monocyte subset, showed prominent enrichment in wt mice, but was less abundant and showed diminished distribution changes after IMQ treatment in *N2^ΔMy^* mice (Figure 3D).

To analyze the initially defined subsets more precisely we applied a multi-parameter gating strategy to define conventional cell subsets (Figure S1C and Table S1) (Gamrekelashvili et al., 2016).

In response to IMQ, Ly6C^hi^ monocytes in wt mice increased transiently in blood, and this response was not altered in mice with conditional Notch2 loss-of function (Figure 3E). In contrast, while Ly6C^lo^ monocytes robustly increased over time with IMQ treatment in wt mice, their levels in *N2^ΔMy^* mice were lower at baseline (Gamrekelashvili et al., 2016) and remained significantly reduced throughout the whole observation period (Figure 3E and F and S2A and B). At the same time, untreated *N2^ΔMy^* mice showed increased levels of MHC-II^+^ atypical monocytes (Figure 3E and F and S2A and B) (Gamrekelashvili et al., 2016), accompanied by a significant wave of macrophages appearing in blood and spleen at d5 (Figure 3E and F and S2A and B). This was followed by a peak in the DC population at d7 (Figure 3E and F and S2A and B). These latter changes did not occur in bone marrow but were only observed in the periphery (Figure S2C and D). Together, these data suggest that wt Ly6C^hi^ monocytes convert to Ly6C^lo^ monocytes in response to TLR stimulation, while Notch2-deficient Ly6C^hi^ monocytes differentiate into MF and DC, suggesting Notch2 as a master regulator of Ly6C^hi^ monocyte cell fate during systemic inflammation.

### Global gene expression analysis identifies expansion of macrophages in blood of Notch2 deficient mice during acute inflammation

To characterize more broadly the gene expression changes involved in monocyte differentiation during inflammation, we next subjected monocyte subsets from PB of wt and *N2^ΔMy^* mice after Sham or IMQ treatment (Figure S3A) to RNA-sequencing and gene expression analysis. After variance filtering and hierarchical clustering, 600 genes were differentially expressed between 6 experimental groups (Figure 4A).

**Figure 3.**
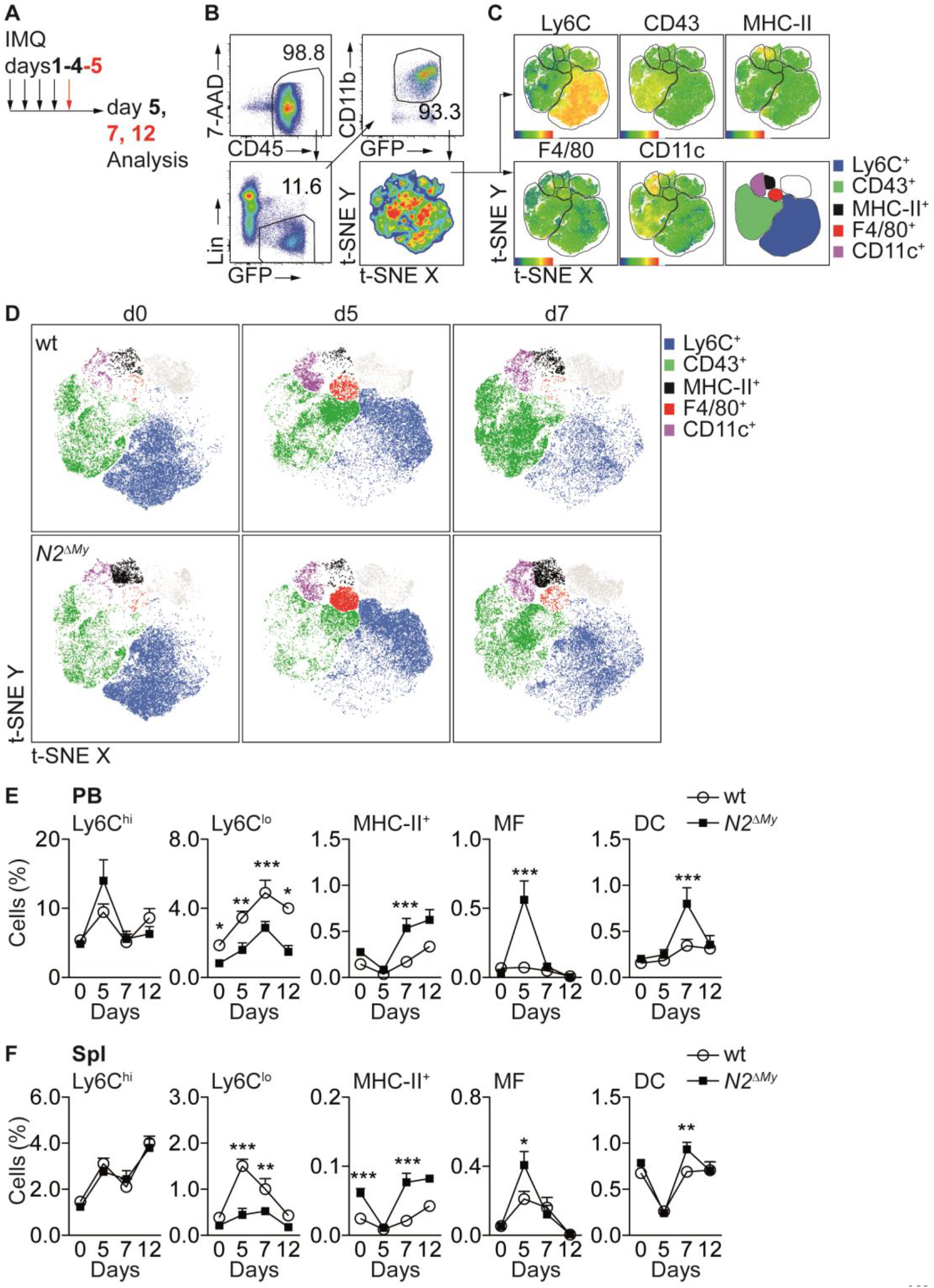
Acute inflammation triggers altered myeloid cell response in *N2^ΔMy^* mice. (**A**) Experimental set-up for IMQ treatment and analysis of mice. (**B, C**) Gating strategy for t-SNE analysis and definition of cell subsets based on expression of surface markers are shown. t-SNE was performed on live CD45^+^Lin^−^CD11b^+^GFP^+^ cells concatenated from 48 PB samples from 4 independent experiments. (**D**) Unsupervised t-SNE analysis showing composition and distribution of cellular subsets from PB of wt or *N2^ΔMy^* IMQ-treated or untreated mice at different time points defined in **B, C** (n=8 mice are pooled for each condition). (**E, F**) Relative frequency of different myeloid subpopulations in PB and Spl of untreated or IMQ-treated mice are shown (data are pooled from 6 experiments n=7-18). (**E, F**) * *P*<0.05, ** *P*<0.01, *** *P*<0.001; 2way ANOVA with Bonferroni’s multiple comparison test.

**Figure 4.**
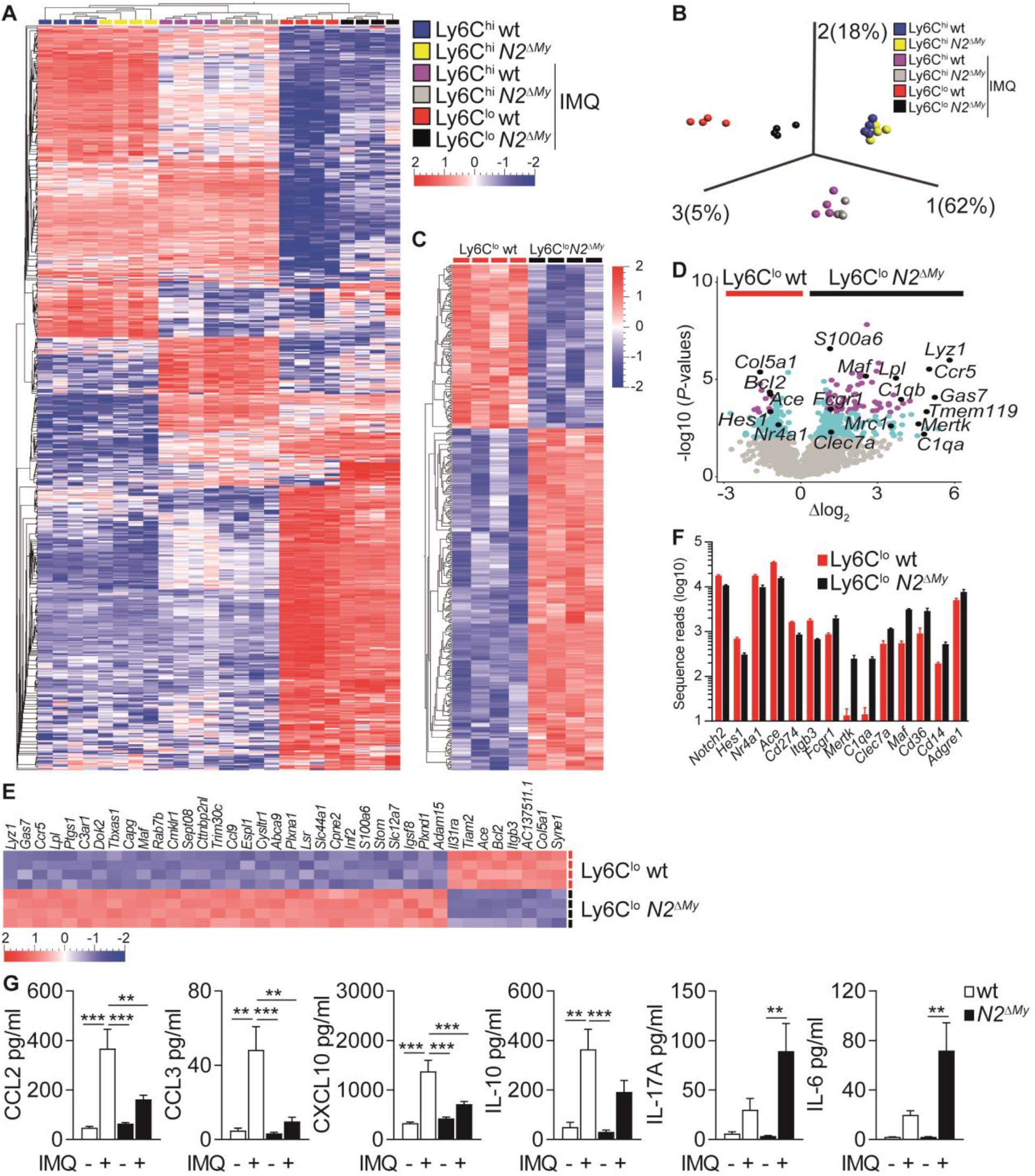
Macrophage development is associated with enhanced inflammatory response in *N2^ΔMy^* mice. (**A, B**) Hierarchical clustering of 600 ANOVA-selected DEGs (**A**) and PCA of PB monocyte subsets (**B**) after IMQ treatment (n=4) is shown (Variance filtering 0.295, ANOVA followed by the B-H correction (*P*<0.0076, FDR≤0.01)). (**C**) Hierarchical clustering of 373 DEGs from IMQ-treated wt and *N2^ΔMy^* Ly6C^lo^ monocyte subsets (Variance filtering 0.117, −0.6≥Δlog_2_≥0.6, Student’s *t*-test with B-H correction (*P*<0.01, FDR≤0.05)). (**D**) Volcano plot showing 379 DEGs between wt and *N2^ΔMy^* LyeC^lo^ cells (−log10(*P*-value)≥2, FDR≤0.05, light blue); and 87 DEGs (−1≥Δlog_2_≥1 (FDR≤0.01) purple); genes of interest are marked black (Student’s *t*-test with B-H correction). (**E**) Heat map of top 38 DEGs from (**D**) log10(*P*-value)≥4, −1≥Δlog_2_≥1, Student’s *t*-test with B-H correction. (**F**) Bar graph showing mean number and SEM of sequence reads for selected genes from IMQ-treated wt and *N2^ΔMy^* Ly6C^lo^ cell subsets. (**G**) Analysis of cytokine and chemokine profiles in the serum of IMQ-treated mice. n=5-10, pooled from 4 independent experiments. (**G**) * *P*<0.05, ** *P*<0.01, *** *P*<0.001; 2way ANOVA with Bonferroni’s multiple comparison test. **Source File1:** List of 600 DEGs for hierarchical clustering and PCA (**Figure 4A, B**) **Source File2:** List of 373 DEGs used for the analysis in **Figure 4C, D** and downstream analysis.

Principal component analysis (PCA) of differentially expressed genes (DEG) of all experimental groups revealed a clear separation between control Ly6C^hi^ monocytes and IMQ-treated Ly6C^hi^ or Ly6C^lo^ monocytes. Interestingly, the effects of Notch2 loss-of-function were most pronounced in the Ly6C^lo^ populations, which separated quite strongly depending on genotype, while Ly6C^hi^ monocytes from wt and *N2^ΔMy^* mice over all maintained close clustering under Sham or IMQ conditions (Figure 4B and C).

Furthermore, while wt Ly6C^lo^ cells were enriched for genes characteristic of patrolling monocytes (*Hes1, Nr4a1, Ace, Cd274 and It9b3),* cells in the Ly6C^lo^ gate from *N2^ΔMy^* mice showed upregulation of genes characteristic of mature phagocytes, such as MF (*Fcgr1, Mertk, C1qa, Clec7a, Maf, Cd36, Cd14, Adgre1* (encoding F4/80)) (Figure 4D-F).

Comparative gene expression analysis of Ly6C^lo^ cell subsets during IMQ treatment identified 373 genes significantly up-or down-regulated with Notch2 loss-of-function (*P*-value <0.01, Figure 4C-F), which were enriched for phagosome formation, complement system components, Th1 and Th2 activation pathways and dendritic cell maturation by ingenuity canonical pathway analysis (Figure S3B). Notably, signatures for autoimmune disease processes were also enriched (Table S2). Independent gene set enrichment analysis (GSEA) (Isakoff et al., 2005; Mootha, Bunkenborg, et al., 2003) confirmed consistent up-regulation of gene sets in *N2^ΔMy^* Ly6C^lo^ cells involved in several gene ontology biological processes, such as vesicle mediated transport (GO:0016192), defense response (GO:0006952), inflammatory response (GO:0006954), response to bacterium (GO:0009617) and endocytosis (GO:0006897) (Table S3). Overall, these data suggest regulation of Ly6C^hi^ monocyte cell fate and inflammatory responses by Notch2.

Furthermore, changes in cell populations resulted in altered systemic inflammatory response patterns. Levels of TLR-induced cytokines and chemokines, such as TNF-α, CXCL1, IL-1β, IFN-α, were elevated to the same extend in wt and *N2^ΔMy^* mutant mice in response to IMQ treatment, suggesting normal primary TLR-activation (Figure S3C). However, circulating levels of chemokines produced by Ly6C^lo^ monocytes (Carlin et al., 2013), such as CCL2, CCL3, CXCL10, and IL-10 were higher in wt mice compared to *N2^ΔMy^* mice, while the levels of pro-inflammatory cytokines IL-17A, IL-6 and GMCSF were strongly enhanced in *N2^ΔMy^* mice, confirming systemic pro-inflammatory alterations in addition to cellular changes in Notch2 loss-of-function mice in response to IMQ (Figure 4G, and S3C).

### Notch2-deficiency promotes macrophage differentiation of Ly6C^hi^ monocytes

To match the observed gene expression pattern of inflammatory Ly6C^lo^ cells from wt and *N2^ΔMy^* mice under IMQ treatment with previously described cells of the monocyte-macrophage lineage we performed pairwise gene set enrichment analysis with defined myeloid cell transcriptomic signatures using the BubbleGUM stand-alone software (Isakoff et al., 2005; Mootha, Bunkenborg, et al., 2003; Spinelli, Carpentier, Montanana Sanchis, Dalod, & Vu Manh, 2015) (Figure 5A). Out of 29 transcriptomic signatures – representing tissue MF, monocyte derived DC (MoDC), conventional DC (cDC), plasmacytoid DC (pDC), classical (Ly6C^hi^) monocytes (cMonocyte), non-classical (Ly6C^lo^ monocytes (ncMonocyte) and B cells (as a reference) – significant enrichment was registered in seven signatures (normalized enrichment score (NES>1.5, FDR<0.1)). The cell fingerprint representing ncMonocyte (#3) was highly enriched in the gene set from wt Ly6C^lo^ monocytes (NES=1.91, FDR=0.039), while all other cell fingerprints showed no significant similarity (Figure 5A and Table S4), confirming a strong developmental restriction towards Ly6C^lo^ monocytes in wt cells. In contrast, *N2^ΔMy^* gene sets showed the highest similarity (NES>1.8 and FDR<0.01) with four cell fingerprints (#1, 5, 6, 7) representing different MF populations, and weak similarity to MoDC (NES=1.64, FDR<0.1) and cMonocyte (NES=1.54, FDR<0.1) (Figure 5A). Phenotyping of cell populations by flow cytometry, using MF markers MerTK and CD64, confirmed selective expansion of MF populations in IMQ-treated *N2^ΔMy^* mice resembling to Lyve-1^hi^MHC-II^lo^-like macrophages, typically associated with blood vessels (Figure 5B and C). Together, these data demonstrate a cell fate switch from Ly6C^lo^ monocytes to macrophages in the absence of Notch2.

**Figure 5.**
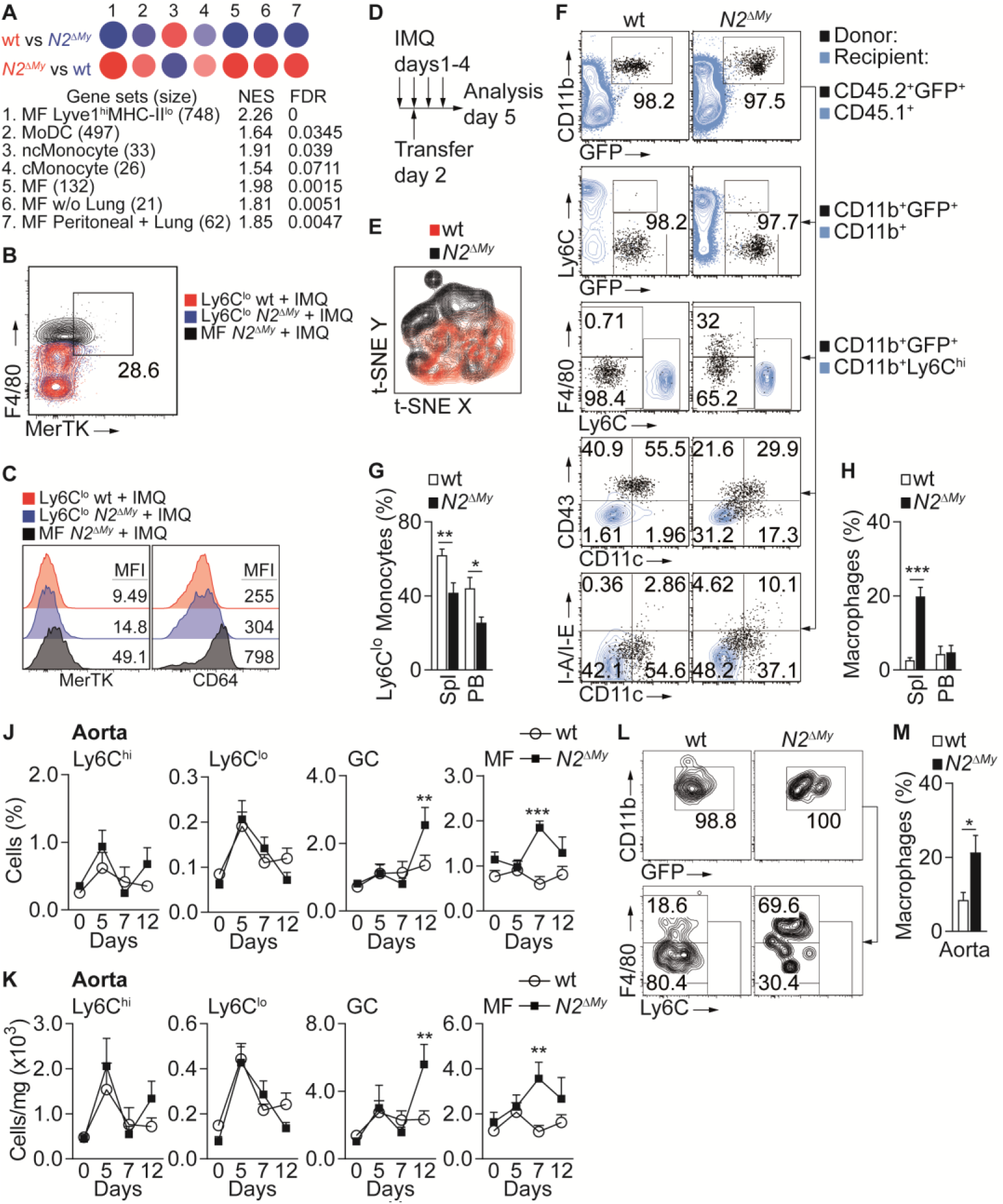
Notch2 deficient Ly6C^hi^ monocytes give rise to macrophages during acute inflammation. (**A**) GSEA based on 373 DEGs between IMQ-treated wt and *N2^ΔMy^* Ly6C^lo^ subsets in PB. Red – positive-, and blue – negative enrichment in corresponding color-coded wt or *N2^ΔMy^* cells. Size of the circle corresponds to NES and intensity of the color to FDR. (**B**) Representative flow cytometry plot showing MerTK expression on Lin-CD11b^+^GFP^+^Ly6C^lo/-^F4/80^hi^ MFs in PB of IMQ-treated *N2^ΔMy^* mice. As a comparison IMQ-treated wt or *N2^ΔMy^* Ly6C^lo^ monocytes are depicted. (**C**) Representative histograms showing expression of surface markers and mean fluorescence intensity (MFI) on myeloid subsets in PB. (**D**) Experimental set-up for IMQ treatment, adoptive transfer and analysis of mice is depicted. (**E**) t-SNE analysis of donor CD45.2^+^CD11b^+^GFP^+^ cells extracted from Spl of IMQ-treated mice. Overlay of wt and *N2^ΔMy^* cells are depicted from concatenated samples. (**F**) Flow cytometry analysis is depicted 3 days after adoptive transfer of wt or *N2^ΔMy^* BM CD45.2^+^ Ly6C^hi^ monocytes in IMQ-treated CD45.1^+^ recipients. Transferred cells are shown in black and recipient CD45.1^+^ (1^st^ row), CD45.1^+^CD11b^+^ (2^nd^ row) or CD45.1^+^CD11b^+^Ly6C^hi^ cells (3^rd^ −5^th^ rows) are depicted in blue. (**G, H**) Frequency of donor-derived Ly6C^lo^ monocytes (**G**) or macrophages (**H**) in CD45.2^+^CD11b^+^GFP^+^ donor cells after adoptive transfer of wt or *N2^ΔMy^* Ly6C^hi^ monocytes is shown. Data are pooled from three independent experiments (n=6/9). (**J, K**) Relative (**J**) and absolute number (**K**) of myeloid subpopulations in aortas of untreated or IMQ-treated mice. Data are pooled from 6 experiments (n=7-18). (**L, M**) Representative Flow cytometry plot (**L**) and frequency of donor-derived macrophages (**M**) in CD45.2^+^CD11b^+^GFP^+^ cells recovered from aortas after adoptive transfer of wt or *N2^ΔMy^* Ly6C^hi^ monocytes. (**M**) Data are pooled from three independent experiments (n=9). (**G, H, M**) * *P*<0.05, ** *P*<0.01, *** *P*<0.001; Student’s *t*-test. (**J, K**) * *P*<0.05, ** *P*<0.01, *** *P*<0.001; 2way ANOVA with Bonferroni’s multiple comparison test. **Source File2:** List of 373 DEGs used for the analysis in **Figure 5A**.

### Notch2 regulates cell fate decisions of Ly6C^hi^ monocytes during inflammation

In the steady-state, Ly6C^hi^ monocytes differentiate into Ly6C^lo^ monocytes and this process is regulated by Notch2 (Gamrekelashvili et al., 2016). In order to confirm that Notch2 controls differentiation potential of Ly6C^hi^ monocytes in response to TLR stimulation we performed adoptive transfer of CD45.2^+^ wt or *N2^ΔMy^* BM Ly6C^hi^ monocytes into IMQ-treated CD45.1^+^ congenic recipients and analyzed the fate of donor cells after three days (Figure 5D). Unsupervised t-SNE analysis of flow cytometry data showed an expanded spectrum of expression patterns in cells from *N2^ΔMy^* donors compared to wt controls (Figure 5E). More precisely, Ly6C^hi^ monocytes from wt mice converted preferentially to Ly6C^lo^ monocytes (Ly6C^lo^F4/80^lo/-^CD11c^+^CD43^+^MHC-II^lo/-^ phenotype) during IMQ treatment (Figure 5F and G). In contrast, conversion of Notch2-deficient Ly6C^hi^ to Ly6C^lo^ monocytes was strongly impaired, but the development of donor-derived F4/80^hi^ macrophages was strongly enhanced (Figure 5H). Furthermore, this expansion of macrophages was also observed in aortas of Notch2-mutant mice *in vivo,* which was followed by granulocyte (GC) infiltration in aortas of IMQ-treated *N2^ΔMy^* mice (Figure 5J and K). Adoptive transfer studies confirmed that MF in IMQ-treated aortas originated from *N2^ΔMy^* Ly6C^hi^ monocytes (Figure 5L and M). These data confirm that Notch2 is a master regulator of Ly6C^hi^ monocyte differentiation potential, regulating a switch between Ly6C^lo^ monocyte or macrophage cell fate during inflammation. These data also demonstrate that in the context of inactive myeloid Notch2 signaling, TLR-stimulation results in systemic pro-inflammatory changes and vascular inflammation.

## Discussion

Together, our data present a spectrum of developmental cell fates of Ly6C^hi^ monocytes and their coordinated regulation by TLR and Notch signaling during inflammation. TLR and Notch signaling act independently and synergistically in promoting Ly6C^lo^ monocyte development from Ly6C^hi^ monocytes, while impairment in either signaling axis impairs Ly6C^lo^ monocyte development. On the other hand, TLR stimulation in the absence of functional Notch2 signaling promotes Ly6C^hi^ monocyte differentiation into MF, suggesting Notch2 as a master regulator of Ly6C^hi^ monocyte cell fate during systemic inflammation.

Plasticity of Ly6C^hi^ monocytes ensures adaptation to environmental signals, which trigger distinct cell developmental programs inducing context- or tissue-specific subsets of terminally differentiated phagocytes, including Ly6C^lo^ monocytes, MF or DC (Guilliams, Mildner, & Yona, 2018). In the steady-state, a subset of Ly6C^hi^ monocytes converts to Ly6C^lo^ monocytes in mice and humans, which is regulated by Notch2 and the endothelial Notch ligand Delta-like 1 (Dll1) (Gamrekelashvili et al., 2016; Patel et al., 2017; Yona et al., 2013). However, when recruited into tissues, Ly6C^hi^ monocytes can give rise to two types of monocyte-derived resident tissue macrophages (MRTM) (Chakarov et al., 2019). These Lyve1^hi^MHC-II^b^ and Lyve1^lo^MHC-II^hi^ MRTMs differ in phenotype and function as well as spatial distribution, but are not typically present in the blood stream. Our gene set enrichment analysis revealed that the signature of cells from inflamed wt mice showed highest and selective similarity to non-classical Ly6C^lo^ monocytes, while Notch2 loss-of-function cells showed the highest similarity to the gene set of Lyve1^hi^MHC-II^lo^ MRTM, but also a more general similarity to an extended spectrum of different MF signatures, suggesting Notch2 as a gate keeper of Ly6C^lo^ monocytes vs. macrophage differentiation. Lyve1^hi^MHC-II^lo^ MFs are closely associated with blood vessels and mediate inflammatory reactions (Chakarov et al 2019). In line with this, Notch2 mutant mice showed elevated numbers of MF in the aorta, but also in the blood stream, and adoptive transfer studies of Ly6C^hi^ monocytes successfully recapitulated their differentiation into aortic MF. These data also have implications for the potential developmental regulation of RTMs by TLR and Notch2. At the same time, the Notch2-mutant population showed weaker but significant enrichment of MoDC signatures, suggesting a mixture of cell subsets representing different stages of monocyte differentiation (Menezes et al., 2016; Mildner et al., 2017) or lineage commitment (Liu et al., 2019; Yanez et al., 2017) within this cell pool. Our genetic targeting strategy acts at or below the level of Ly6C^hi^ monocytes (Gamrekelashvili et al., 2016) and Ly6C^hi^ monocyte numbers in Notch2-mutant mice are not altered at steady-state or during TLR stimulation, which suggests cell fate decision at or below the level of Ly6C^hi^ monocytes. However, whether Notch2 loss-of-function differentially impacts monocyte subset composition requires further study.

While our current data clearly demonstrate that Notch2 loss-of-function promotes macrophage development from Ly6C^hi^ monocytes and a pro-inflammatory milieu during TLR stimulation, we have previously shown that Dll1-Notch signaling promotes maturation of anti-inflammatory macrophages from Ly6C^hi^ monocytes in ischemic muscle (Krishnasamy et al., 2017). Furthermore, Dll4-Notch signaling initiated in the liver niche was recently shown to promote Kupffer cell development after injury (Johnny Bonnardel et al., 2019; Sakai et al., 2019) or to promote pro-inflammatory macrophage development (Xu et al., 2012). This suggests that the role of Notch is ligand-, cell- and context-specific, which emphasizes the differential effects of specific ligand-receptor combinations (Benedito et al., 2009). Our data demonstrate that Notch2 is a master regulator of Ly6C^hi^ monocyte cell fate during inflammation, which contributes to the nature of the inflammatory response.

Lastly, our data also reveal in important function of myeloid Notch2 for regulation of systemic and vascular inflammation with potential implications for autoimmune disease. When wt mice are challenged with TLR stimulation they show predominant conversion of Ly6C^hi^ monocytes into Ly6C^lo^ monocytes with blood vessel patrolling and repairing function (Carlin et al., 2013), IL-10 secretion, but no evidence of vascular inflammation. In contrast, Notch2 mutant mice show predominant and ectopic differentiation of Ly6C^hi^ monocytes into macrophages, which appear in the blood stream and the spleen and infiltrate major blood vessels, such as the aorta, along with systemic increases in pro-inflammatory cytokines. In addition, absence of functional Notch2 promoted a core macrophage signature and strong upregulation of canonical pathways involved in autoimmune disease. Since TLR7 has been implicated in the development of autoimmune disease (M. L. Santiago-Raber et al., 2010), our data suggest Notch2 as an important modulator of this process, with active Notch2 protecting from overt inflammation, but inactive Notch2 promoting overt inflammation.

## Materials and methods

### Mice

*Cx3cr1^GFP/+^* mice(Jung et al., 2000), *LysM^Cre^* mice(Clausen, Burkhardt, Reith, Renkawitz, & Forster, 1999), *Notch2^lox/lox^* mice (Besseyrias et al., 2007), *Cx3cr1^GFP/+^LysM^Cre^Notch2^lox/lox^* (*N2^ΔMy^*) (Gamrekelashvili et al., 2016) have been previously described. B6.SJL-*Ptprc^a^Pepc^b^*/BoyJ (CD45.1^+^) mice were purchased from central animal facility of Hannover Medical School (ZTL, MHH). *B6.129P2-Myd88^tm1Hlz^/J* (*Myd88^−/−^*) (Gais et al., 2012) and *Myd88^+/+^* littermate control (wt) mice were kindly provided by Dr. Matthias Lochner. Mice were housed under specific pathogen free conditions in the animal facility of Hannover Medical School. All experiments were performed with 8-12 weeks old mice and age and sex matched littermate controls with approval of the local animal welfare board (LAVES, Lower Saxony, Animal Studies Committee).

### Tissue and cell preparation

For single cell suspension mice were sacrificed and spleen, bone marrow, blood and aortas were collected. Erythrocytes were removed by red blood cell lysis buffer (BioLegend) or by density gradient centrifugation using Histopaque 1083 (Sigma-Aldrich). Aortas were digested in DMEM medium supplemented with 500U/ml Collagenase II (Worthington). After extensive washing cells were resuspended in PBS containing 10%FCS and 2mM EDTA kept on ice, stained and used for flow cytometry or for sorting.

### Flow cytometry and cell sorting

Non-specific binding of antibodies to Fc-receptors was blocked with anti-mouse CD16/CD32 (TruStain fcX from BioLegend) in single cell suspensions prepared from Spl, PB or BM. After subsequent washing step cells were labelled with primary and secondary antibodies or streptavidin-fluorochrome conjugates and were used for flow cytometry analysis (LSR-II, BD Biosciences) or sorting (FACSAria; BD Biosciences or MoFlo XDP; Beckman Coulter). Antibodies and fluorochromes used for flow cytometry are described in Table S5. Flow cytometry data were analyzed using FlowJo software (FlowJo LLC). Initially cells were identified based on FSC and SSC characteristics. After exclusion of doublets (on the basis of SSC-W, SSC-A), relative frequency of each subpopulation from live cell gate, or absolute number of each subset (calculated from live cell gate and normalized per mg BM, mg Spl, mg aorta or μl PB) were determined and are shown in the graphs as mean ± SEM, unless otherwise stated. Unsupervised t-distributed stochastic neighbor embedding (t-SNE) analysis (van der Maaten & Hinton, 2008) was performed on live CD45^+^Lin^−^GFP^+^CD11b^+^ population in concatenated samples using FlowJo.

### Cytokine multiplex bead-based assay

Sera were collected from control or Aldara treated mice and kept frozen at −80°C. Concentration of IFN-γ, CXCL1, TNF-α, CCL2, IL-12(p70), CCL5, IL-1β, CXCL10, GM-CSF, IL-10, IFN-β, IFN-α, IL-6, IL-1α, IL-23, CCL3, IL-17A were measured with LEGENDplex multi-analyte flow assay kits (BioLegend) according to manufacturer’s protocol on LSR-II flow cytometer. Data were processed and analyzed with LEGENDplex data analysis software (BioLegend).

### RNA isolation, library construction, sequencing and analysis

Peripheral blood monocyte subpopulations were sorted from Aldara treated mice or untreated controls (Figure S3A) and RNA was isolated using RNeasy micro kit (Qiagen). Libraries were constructed from total RNA with the ‘SMARTer Stranded Total RNA-Seq Kit v2 – Pico Input Mammalian’ (Takara/Clontech) according to manufacturer’s recommendations, barcoded by dual indexing approach and amplified with 11 cycles of PCR. Fragment length distribution was monitored using ‘Bioanalyzer High Sensitivity DNA Assay’ (Agilent Technologies) and quantified by ‘Qubit^®^ dsDNA HS Assay Kit’ (ThermoFisher Scientific). Equal molar amounts of libraries were pooled, denatured with NaOH, diluted to 1.5pM (according to the Denature and Dilute Libraries Guide (Illumina)) and loaded on an Illumina NextSeq 550 sequencer for sequencing using a High Output Flowcell for 75bp single reads (Illumina). Obtained BCL files were converted to FASTQ files with bcl2fastq Conversion Software version v2.20.0.422 (Illumina). Pre-processing of FASTQ inputs, alignment of the reads and quality control was conducted by nfcore/rnaseq (version 1.5dev) analysis pipeline (The National Genomics Infrastructure, SciLifeLab Stockholm, Sweden) using Nextflow tool. The genome reference and annotation data were taken from GENCODE.org (GRCm38.p6 release M17). Data were normalized with DESeq2 (Galaxy Tool Version 2.11.39) with default settings and output counts were used for further analysis with Qlucore Omics explorer (Sweden). Data were log_2_ transformed, 1.1 was used as a threshold and low expression genes (<50 reads in all samples) were removed from the analysis. Hierarchical clustering (HC) or principal component analysis (PCA) was performed on 600 differential expressed genes (DEGs) after variance filtering (filtering threshold 0.295) selected by ANOVA with the Benjamini-Hochberg (B-H) correction (*P*<0.01, FDR≤0.01). For two group comparisons Student’s *t*-test with B-H correction was used and 373 DEGs ((Variance filtering 0.117, −0.6≥Δlog_2_≥0.6 (*P*<0.01, FDR≤0.05)) were selected for further IPA or GSEA analysis.

Ingenuity pathway analysis (IPA) was performed on 373 DEGs using IPA software (Qiagen) with default parameters. Top 20 canonical pathways and top 5 immunological diseases enriched in DEGs were selected for display.

Gene set enrichment analysis (GSEA) (Mootha, Lindgren, et al., 2003; Subramanian et al., 2005) was performed on 373 DEGs using GSEA software (Broad institute) and C5 GO biological process gene sets (Liberzon et al., 2015) from MsigDB with 1000 gene set permutations for computing *P*-values and FDR.

BubbleGUM software, an extension of GSEA (Broad Institute) (Spinelli et al., 2015; Vu Manh, Elhmouzi-Younes, et al., 2015), and published transcriptomic signatures (Chakarov et al., 2019; Gautier et al., 2012; Schlitzer et al., 2015; Vu Manh, Bertho, Hosmalin, Schwartz-Cornil, & Dalod, 2015) were used to assess and visualize the enrichment of obtained gene sets for myeloid populations and define the nature of the cells from which the transcriptomes were generated.

### *In vitro* conversion studies

96 well flat bottom plates were coated at room temperature for 3 hours with IgG-Fc or DLL1-Fc ligands (all from R&D) reconstituted in PBS. Sorted BM Ly6C^hi^ monocytes <4 were cultured in coated plates and were stimulated with Resiquimod (R848, 0.2μg ml^−1^, Cayman Chemicals) or LPS (0.2μg ml^−1^, E. coli O55:B5 Sigma-Aldrich) in the presence of M-CSF (10ng ml^−1^, Peprotech) at 37 °C for 24 hours. One day after culture, cells were harvested, stained and subjected to flow cytometry. Frequency of Ly6C^lo^ monocyte-like cells (CD11b^+^GFP^+^Ly6C^lo/-^CD11c^lo^MHC-II^lo/-^CD43^+^) in total live CD11b^+^GFP^+^ cells served as an indicator of conversion efficiency and is shown in the graphs. Alternatively, cultured cells were harvested and isolated RNA was used for gene expression analysis.

### Induction of acute systemic inflammation using Aldara

Mice were anesthetized and back skin was shaved and depilated using depilating crème. Two days after 50mg/mouse/day Aldara (containing 5% Imiquimod, from Meda) or Sham crème were applied on depilated skin and right ear (where indicated) for 4-5 consecutive days (El Malki et al., 2013; van der Fits et al., 2009). Mouse weight and ear thickness were monitored daily. Mice were euthanized on the indicated time points after start of treatment (day 0, 5, 7 and 12) PB, Spl, BM and aortas were collected for further analysis.

### Adoptive cell transfer experiments

CD11b^+^Ly6C^hi^GFP^+^ monocytes were sorted from BM of CD45.2^+^ donors and injected into CD45.1^+^ recipients intravenously (i.v.). In separate experiment wt or *Myd88^−/−^* CD45.2^+^CD11b^+^Ly6C^hi^CXsCR1^+^ monocytes were used as a donor for transfer. 30 minutes later PBS or R848 (37.5ug per mouse) were injected in recipient mice. Two days after transfer Spl, PB and BM were collected and single cell suspension was prepared. After blocking of Fc receptors (anti-mouse CD16/CD32, TruStain fcX from BioLegend), cells were labelled with biotin-conjugated antibody cocktail containing anti-CD45.1 and anti-Lin (anti-CD3, CD19, B220, NK1.1, Ly6G, Ter119) antibodies, antibiotin magnetic beads and enriched on LS columns (Miltenyi Biotec) according to manufacturer’s instructions. CD45.1^neg^Lin^neg^ fraction was collected, stained and analyzed by flow cytometry. Ly6C^lo^ monocytes (CD45.2^+^CD11b^+^GFP^+^Ly6C^lo/-^ F4/80^lo^CD11c^lo^CD43^hi^ cells) and macrophages (CD45.2^+^CD11b^+^GFP^+^Ly6C^lo/-^F4/80^hi^ cells) were quantified in Spl, BM and PB as relative frequency of total donor derived CD45.2^+^CD11b^+^GFP^+^ cells. Adoptive transfer experiments in IMQ-treated mice were performed using wt or *N2^ΔMy^* CD11b^+^Ly6C^hi^GFP^+^ donor monocytes and Spl, PB and aortas of recipients were analyzed 3 days after transfer.

### Quantitative real-time PCR analysis

Total RNA was purified from cell lysates using Nucleospin RNA II kit (Macherey Nagel). After purity and quality check, RNA was transcribed into cDNA employing cDNA synthesis kit (Invitrogen) according to manufacturer’s instructions. Quantitative realtime PCR was performed using specific primers for *Nr4a1*: forward, 5’-AGCTTGGGTGTTGATGTTCC-3’, reverse, 5’-AATGCGATTCTGCAGCTCTT-3’ and *Pou2f2:* forward, 5’-TGCACATGGAGAAGGAAGTG-3’, reverse, 5’-AGCTTGGGACAATGGTAAGG-3’ and FastStart Essential DNA Green Master on a LightCycler 96 system from Roche according to manufacturer’s instructions. Expression of each specific gene was normalized to expression of *Rps9* and calculated by the comparative CT (2^−ΔΔCT^) method (Schmittgen & Livak, 2008).

### Statistical analysis

Results are expressed as mean ± standard error of mean (SEM). N numbers are biological replicates of experiments performed at least three times unless otherwise indicated. Significance of differences was calculated using unpaired, two-tailed Student’s *t-*test with confidence interval of 95%. For comparison of multiple experimental groups one-way or two-way ANOVA and Bonferroni’s multiple-comparison test was performed.

### Data availability

Data from RNA sequencing have been deposited to NCBI’s Gene Expression Omnibus and are available under the accession number GSE147492.

All the relevant data are available upon request.

## Acknowledgements

We thank the Central Animal Facility, Research Core Facility Cell Sorting and Research Core Unit Genomics of Hannover Medical School for support. We thank Thien-Phong Vu Manh and Lionel Spinelli for consultation on BubbleGUM.

Funded by grants from DFG (GA 2443/2-1), DSHF (F/17/16) to J.G., and DFG (Li948-7/1) to FPL.

## Author contributions

J.G., T.K., S.S., C.R., K.D., A.S., S.F. did the experiments;

J.G., C.R., F.P.L. designed experiments and analyzed data;

S.K., P.W., M.L., B.H., T.S., U.K., H.H. provided necessary materials or animals;

J.G. and F.P.L. wrote and edited the manuscript;

All authors corrected the manuscript;

J.G. and F.P.L. conceived and directed the study.

## Supplementary Information

**Figure S1.**
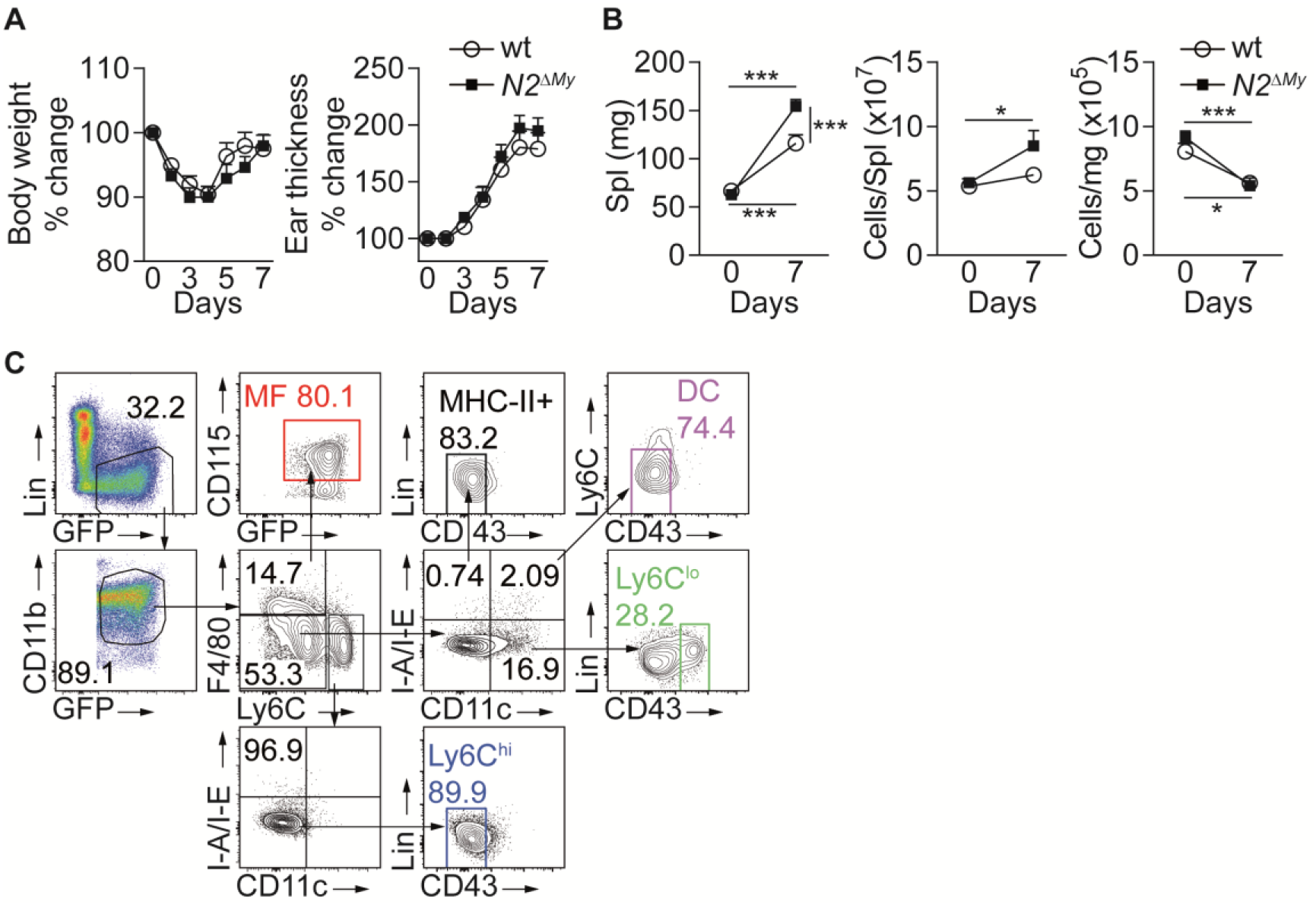
IMQ-mediated inflammation and gating strategy for myeloid cell identification. (**A, B**) IMQ treatment induces systemic inflammation in mice: (**A**) Body weight (left) and ear thickness (right) of IMQ-treated wt or *N2^ΔMy^* mice (data are from 3 experiments, n=11/13). (**B**) Spleen weight and cell number of IMQ-treated wt or *N2^ΔMy^* mice (data are from 3 independent experiments, n=11/12). (**C**) Gating strategy for definition of myeloid subsets in flow cytometry analysis. (**B**) * *P*<0.05, *** *P*<0.001; 2way ANOVA with Bonferroni’s multiple comparison test.

**Figure S2.**
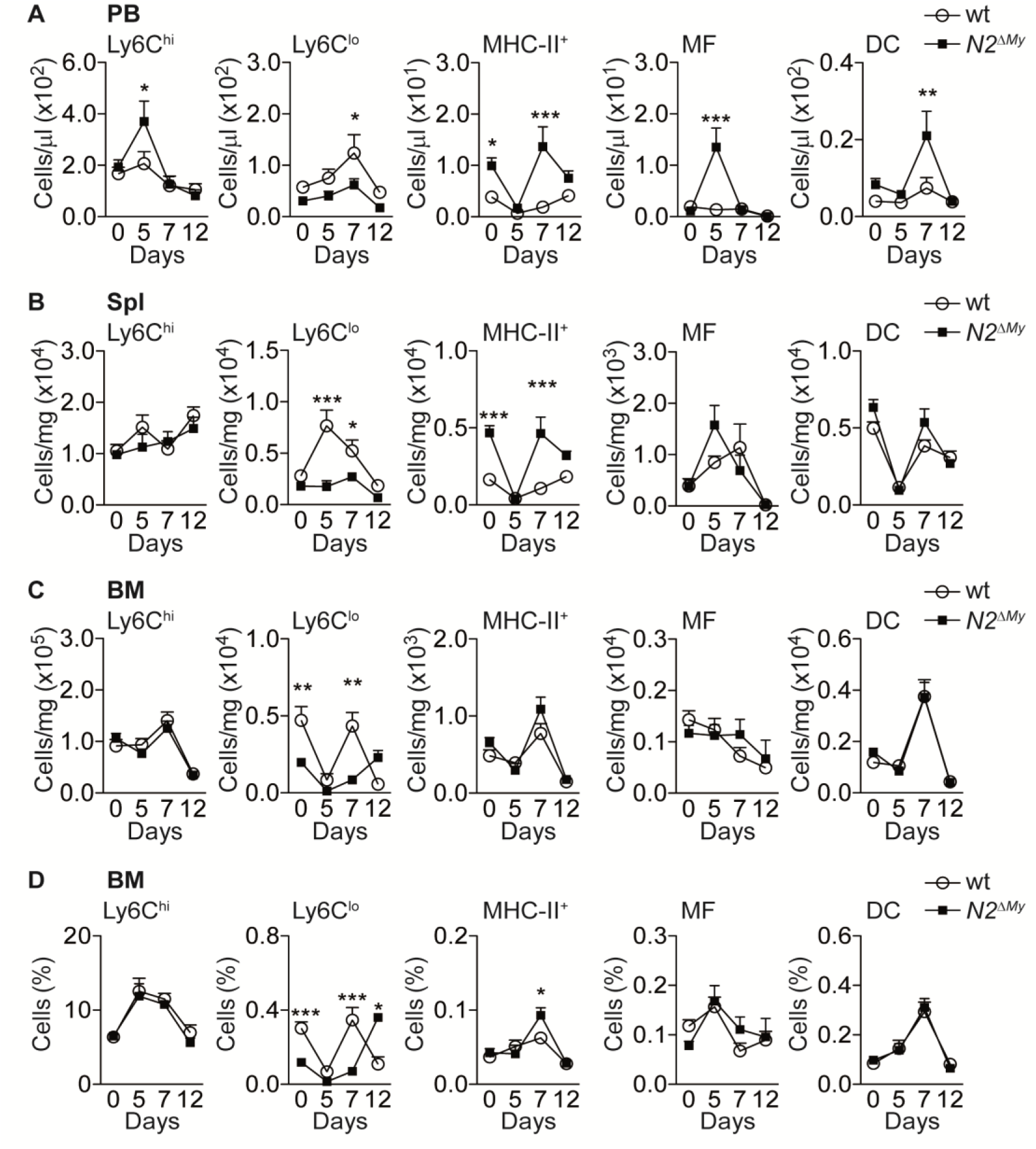
Flow cytometry analysis of myeloid cells in IMQ-driven inflammation. (**A-D**) Relative and absolute frequency of different myeloid subpopulations in PB, spl or BM of IMQ-treated mice from 6 experiments are shown (n=7-18). * *P*<0.05, ** *P*<0.01, *** *P*<0.001; 2way ANOVA with Bonferroni’s multiple comparison test.

**Figure S3.**
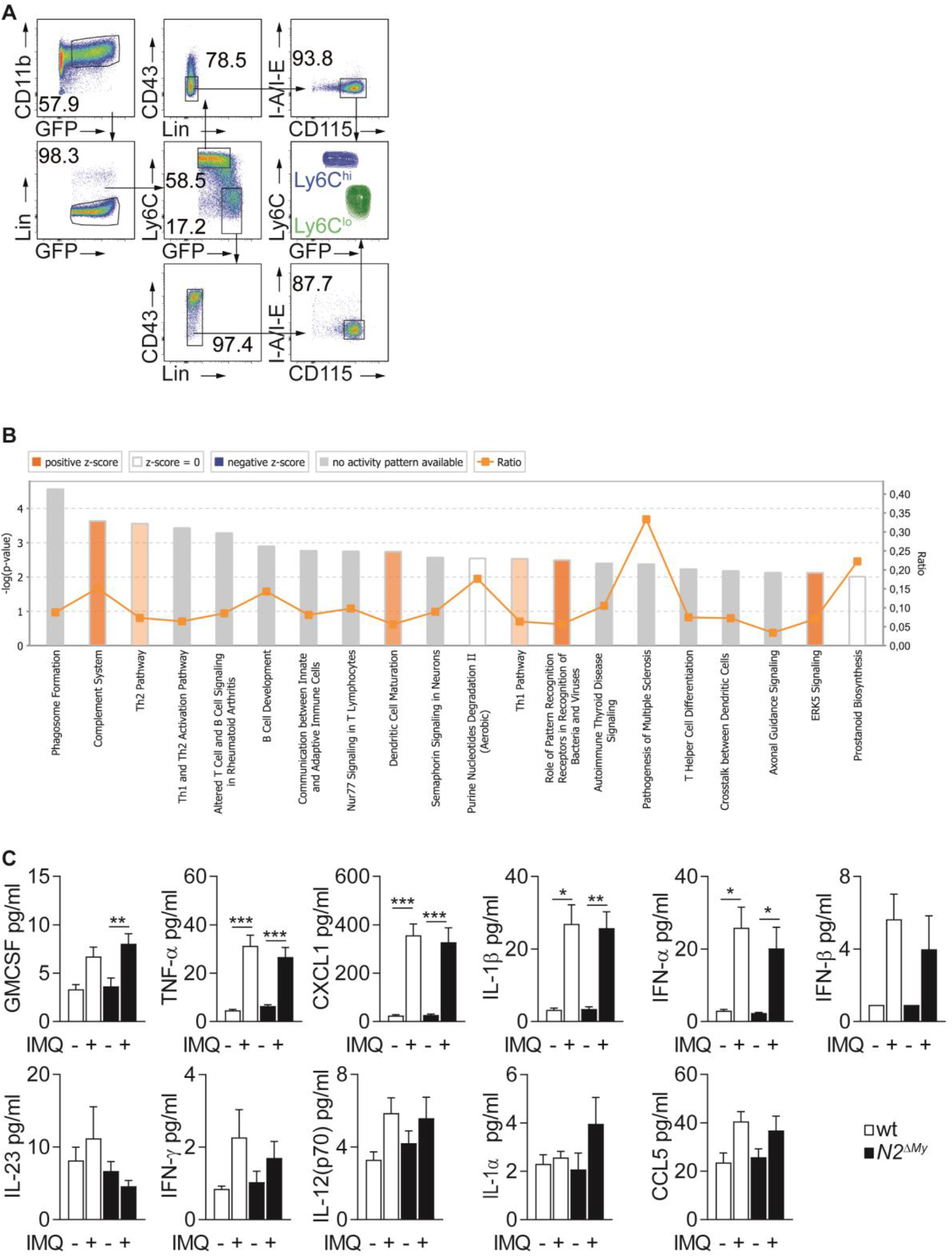
Gating strategy for cell sorting, IPA and Cytokine and chemokine analysis in IMQ-treated mice. (**A**) Sorting strategy of monocyte subsets for RNA-seq analysis. Lin^−^ CD11b^+^GFP^+^CD115^+^Ly6C^hi^CD43^−^MHC-II^−^ (Ly6C^hi^) and Lin^−^CD11b^+^GFP^+^CD115^+^Ly6C^lo/-^MHC-II^−^ (Ly6C^lo^) monocytes were sorted from naïve or IMQ-treated wt or *N2^ΔMy^* mouse PB and used for RNA-seq. (**B**) top 20 Ingenuity canonical pathways enriched in mutant Ly6C^lo^ cells as compared to wt Ly6C^lo^ subset from IMQ-treated mice. (**C**) Analysis of chemokine and cytokine levels in untreated or IMQ-treated wt or *N2^ΔMy^* mouse serum. Data are pooled from 4 independent experiments (n=5-10).* *P*<0.05, ** *P*<0.01, *** *P*<0.001; 2way ANOVA, Bonferroni’s post-test. **Source File2:** List of 373 DEGs used for the analysis in **Figure S3B**.

## Supplementary Tables

**Table S1.**
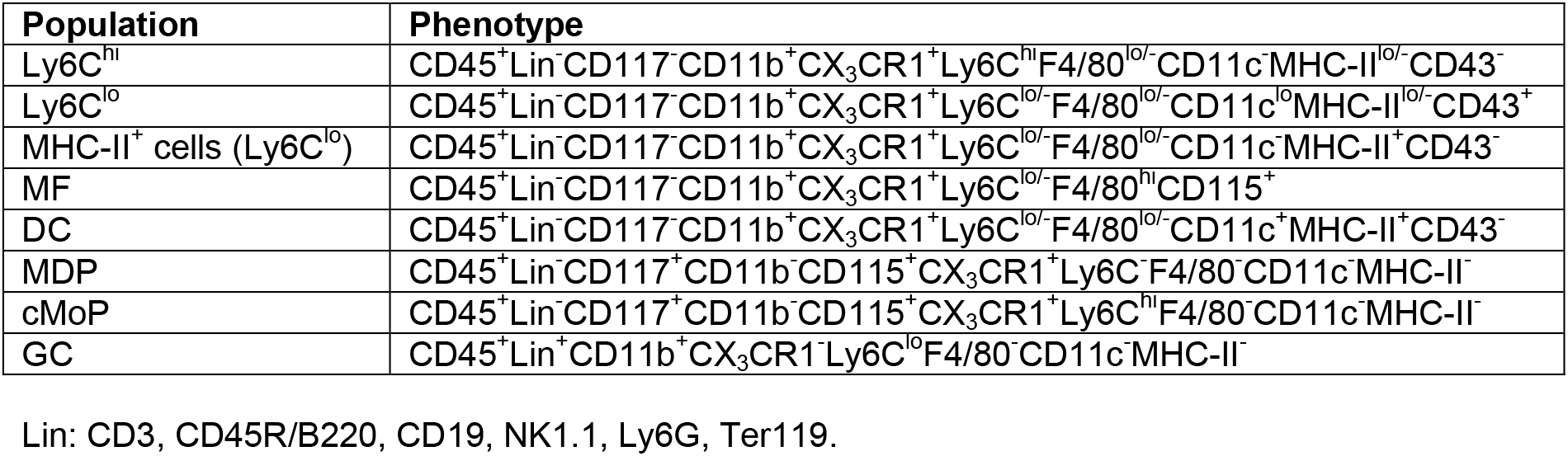
Surface phenotype signatures for identification of distinct myeloid populations *in vivo.*

**Table S2.**
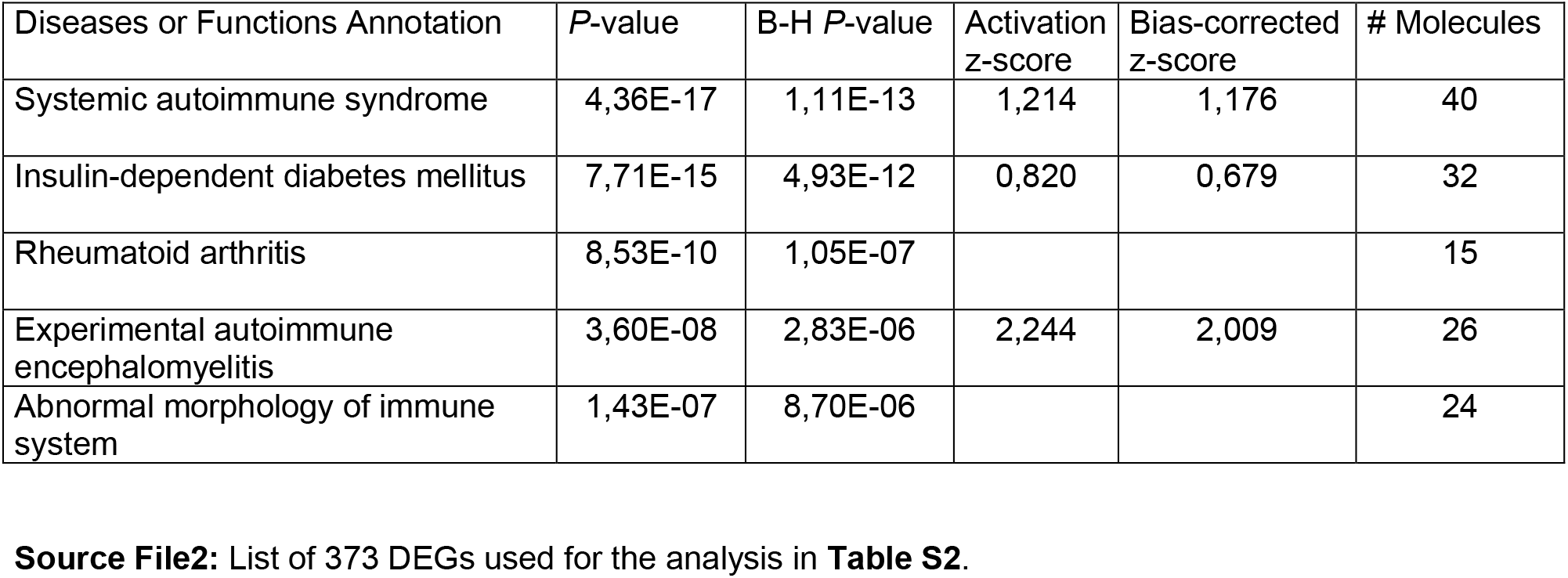
IPA of top 5 immunological diseases enriched in Ly6C^lG^ cells from IMQ-treated *N2^ΔMy^* mice.

**Table S3.**
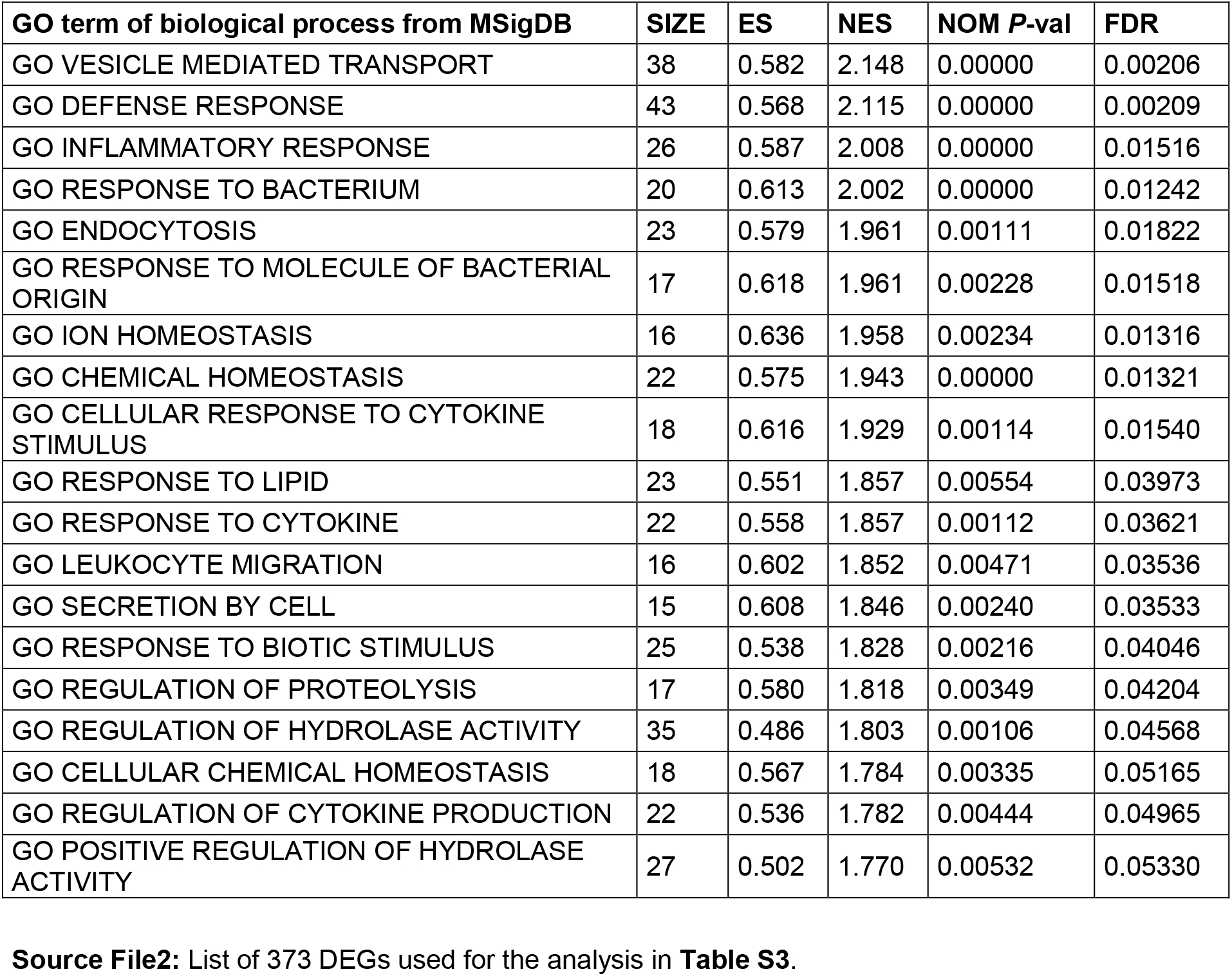
Top 20 gene sets involved in GO biological processes enriched in mutant Ly6C^lo^ monocytes from IMQ-treated mice.

**Table S5.** Antibodies and fluorescence dyes for flow cytometry used in the study.

| **Antibody/Dye** | **Clone** | **Company** |
| --- | --- | --- |
| CD3 | 17A2 | BioLegend |
| Ter119 | Ter119 | BioLegend |
| CD45R/B220 | RA3-6B2 | BioLegend |
| Ly6G | 1A8 | BioLegend |
| CD19 | 6D5 | BioLegend |
| NK1.1 | PK136 | BioLegend |
| CD117 | 2B8 | BioLegend |
| CD115 | AFS98 | BioLegend |
| CD11b | M1/70 | BioLegend |
| Ly6C | HK1.4 | BioLegend |
| F4/80 | BM8 | BioLegend |
| CD11c | N418 | BioLegend |
| I-A/I-E | M5/114.15.2 | BioLegend |
| CD64 | X54-5/7.1 | BioLegend |
| CD45 | 30-F11 | BioLegend |
| CD45.1 | A20 | BioLegend |
| CD45.2 | 104 | BioLegend |
| CX<sub>3</sub>CR1 | SA011F11 | BioLegend |
| MerTK | DS5MMER | eBioscience |
| CD43 | S7 | BD Pharmingen |
| Streptavidin-PE-Dazzled594 |  | BioLegend |
| 7AAD |  | BioLegend |
| Propidium Iodide |  | Sigma |

